# Implementation of single molecule FRET for visualizing intramolecular movement in CRISPR-Cas9

**DOI:** 10.1101/2020.04.13.039537

**Authors:** Haruka Narita, Hiroshi Ebata, Karibu Sakai, Katsuhiko Minami, Sotaro Uemura, Tomohiro Shima

## Abstract

**SHORT ABSTRACT:** This paper summarizes how to visualize the flexible inter-domain movements of CRISPR-associated protein Cas9 using single molecule FRET

**LONG ABSTRACT:** The CRISPR-associated protein Cas9 is widely used as a genome editing tool because of its ability to be programmed to cleave any DNA sequence that is followed by a protospacer adjacent motif. The continuing expansion of Cas9 technologies has stimulated studies regarding the molecular basis of the Cas9 catalytic process. Here we summarize methods for single molecule FRET (smFRET) to visualize the inter-domain movements of Cas9 protein. Our measurements and analysis demonstrate flexible and reversible movements of the Cas9 domains. Such flexible movements allow Cas9 to adopt transient conformations beyond those solved by crystal structures and play important roles in the Cas9 catalytic process. In addition to the smFRET measurement itself, to obtain precise results, it is necessary to validate Cas9 catalytic activity. Also, fluorescence anisotropy data are required to interpret smFRET data properly. Thus, in this paper, we describe the details of these important additional experiments for smFRET measurements.

## INTRODUCTION

The clustered regularly interspaced short palindromic repeats (CRISPR)-CRISPR-associated proteins (Cas) system was first identified as a prokaryotic immune system that protects prokaryotes from viral attack through acquired immunity^1–5^. Among the various CRISPR-Cas systems, CRISPR-Cas9 employs a single Cas9 protein that can be programmed to cleave any specific DNA (deoxyribonucleic acid) sequence that is followed by a protospacer adjacent motif (PAM) through its binding with single guide ribonucleic acid (sgRNA)^6, 7^. The programmable DNA-cleavage ability of Cas9 leads it to play a central role in the field of genome engineering^8, 9^. Recently, much effort has been devoted to improve the efficacy and specificity of CRISPR-Cas9 gene-editing^10, 11^. Also, the application of Cas9 has been expanded to a wide variety of biological research techniques, such as the visualization and regulation of genomic DNA^12–15^.

Improvements and applications of the CRISPR-Cas9 system have brought greater attention to the molecular basis of the Cas9 catalytic process. Structural studies have revealed a series of conformational changes in Cas9 during the process^16–20^. The Cas9 protein is mainly composed of recognition (REC) and nuclease (NUC) lobes. Upon binding with sgRNA, the two lobes relatively rotate to convert Cas9 into an active conformation that can bind to the target DNA. After binding to the target DNA, the two catalytic domains in the NUC lobe (HNH and RuvC) translocate their positions in the protein and cleave opposite strands of the DNA. These structural studies have deepened our understanding of the catalytic process; however, the dynamics of the conformational changes have remained elusive. In particular, no crystal structures have shown direct interactions of the HNH catalytic domain and the cleavage point in the target DNA^18, 20^. Although bulk FRET measurements have proposed concerted DNA cleavage by HNH and RuvC^21^, the averaged data of bulk conditions provides limited information about the DNA cleavage mechanism. Instead, methods that detect when and how HNH translocates to the cleavage point are required.

Here we explain how single-molecule fluorescence resonance energy transfer (smFRET) measurements can monitor the dynamics of Cas9 conformational changes. smFRET can monitor the dynamics of two fluorescent dyes (donor and acceptor) labeling a target molecule with sub-nanometer accuracy and millisecond time resolution^22^. We have applied this system to reveal the flexible and reversible movements of the HNH domain in the DNA-sgRNA-Cas9 ternary complex^23^. The HNH domain was positioned at the DNA cleavage point only during flexible movements, suggesting the importance of domain flexibility in the Cas9 catalytic process. These results demonstrate that smFRET is well suited for detecting inter-domain flexibility because of its high spatial and temporal resolution but requires careful preparation before the measurement. While general smFRET methodologies have already been introduced by several papers^22, 24^, we describe the preparation and analysis for successful smFRET measurements for Cas9 in detail.

## PROTOCOL

### Fluorescent labeling of Cas9

All three Cas9 constructs for FRET measurement (D435C-E945C, S355C-S867C and S867C-N1054C) contain only two cysteine residues and are fluorescently labeled using Cyanine dye 3- and Cyanine dye 5-maleimide.

1. Preparation of Cyanine dye 3- and Cyanine dye 5-maleimide stock solution

1.1 Dissolve Cyanine dye 3- and Cyanine dye 5-maleimide powder (one vial for labeling 1 mg/mL antibody) separately in 20 μL dimethyl sulfoxide (DMSO) to make approximately 20 mM stock solutions.
1.2 To determine the exact dye concentration, take small aliquot of the stock (1-2 μL) and dilute it 10,000 times by desterilized water. Measure the absorbance of the diluted solution of Cyanine dye 3-maleimide at 550 nm and of Cyanine dye 5-maleimide at 649 nm using an absorption spectrophotometer. The values of measured absorbance should be in the range of between 0.2 and 0.8, in which range the relationship between absorbance and concentration is linear according to the Beer-Lambert law. Otherwise, adjust the dilution ratio and measure absorbance again.
1.3 Calculate the concentration of Cyanine dye 3- and Cyanine dye 5-maleimide in the diluted solution using the molar extinction coefficient of Cyanine dye 3-maleimide (150,000 M^−1^cm^−1^ at 550 nm) and Cyanine dye 5-maleimide (250,000 M^−1^cm^−1^ at 649 nm), respectively.
1.4 Calculate the original concentration in the stock solutions according to the dilution ratio used in step 1.2. Dilute the stock solution with DMSO to make 10 mM stock for future use.
1.5 Aliquot the 10 mM stocks into 0.2-mL PCR (polymerase chain reaction) tubes. To avoid hydrolysis, store the tubes in vacuum-sealed bags at −30 °C until use.
2. Fluorescent labeling of Cas9

2.1 Exchange buffer of the stocked Cas9 solution to labeling buffer (20 mM HEPES (4-(2-hydroxyethyl)-1-piperazineethanesulfonic acid)-KOH pH 7.0, 100 mM KCl, 2 mM MgCl_2_, 5% glycerol) by using a gel filtration spin column for 20 – 75 μL solution (cut off molecular weight of 40 kDa). Follow the manufacturer’s protocol with one modification. NOTE: At the N-terminus of Cas9, BCCP (biotin carboxyl carrier protein)-tag is attached for biotinylation in E. Coli (Escherichia coli) during protein expression. Since BCCP-tag has no cysteine residue, it does not affect maleimide labeling. The influence of BCCP-tag insertion on Cas9 activity was also evaluated using the DNA cleavage assay described below.

2.1.1. Place the column in a centrifuge tube. Centrifuge the column for 2 min in a microcentrifuge at 1,000 × g, 4 °C to remove liquid in the column.
2.1.2. Apply 500 μL of labeling buffer onto the gel resin and centrifuge for 1 min at 1,000 × g, 4 °C to equilibrate the column. Discard liquid that flows through the column. Repeat this equilibration step 3 times.
2.1.3. Apply an equivalent volume of labeling buffer to that of the sample being used in the step 2.1.4. (20-75 μL) onto the resin. Centrifuge for 4 min at 1,000 × g, 4 °C. NOTE: This is the modification step from the manufacturer’s protocol. By including this step, the protein solution can be eluted without diluting the original solution.
2.1.4. Place a column in a clean 1.5 mL microcentrifuge tube. Carefully apply the sample (20-75 μL) to the column.
2.1.5. After loading the sample, centrifuge the column for 4 min, 1,000 × g, 4 °C. Collect the flow-through sample solution.
2.2. Add 0.5 mM Tris(2-carboxyethyl)phosphine hydrochloride (TCEP) to the Cas9 solution and incubate on ice for 30 min to reduce disulfide bonds in Cas9. NOTE: DO NOT use dithiothreitol (DTT) or β-mercaptoethanol to reduce the disulfide bonds, since these compounds contain sulfhydryl groups that react with maleimide.
2.3. Mix Cyanine dye 3- and Cyanine dye 5-maleimide with the Cas9 solution at a 1:20 (protein:dye) molar ratio. NOTE: Fluorescent maleimide specifically reacts with the cysteine sulfhydryl group (Fig. 1).
2.4. Following incubation of the mixed solution on ice for 2 h, quench the reaction by adding 10 mM DTT. NOTE: Since DTT contains a sulfhydryl group, the addition of excessive DTT quenches the activity of maleimide.
2.5. Remove excess fluorescent maleimide dye using a spin column gel filtration with assay buffer (AB: 20 mM HEPES-KOH pH 7.5, 100 mM KCl, 2 mM MgCl_2_, 5% glycerol, 0.5 mM EDTA, 1 mM DTT) following the procedure described in step 2.1. Repeat this step twice for complete removal of excess dye.
2.6. Measure the concentration of the labeled Cas9 and labeling efficiency.

2.6.1. Measure the absorbance of the labeled Cas9 at 280 nm and the absorption peak of Cyanine dye 3 (550 nm) and Cyanine dye 5 (649 nm), respectively. NOTE: Use a clean disposal cuvette with ~50 μL capacity and collect the sample in the cuvette following the measurement.
2.6.2. Calculate the concentration of the labeled Cas9 and the labeling efficiency using the molar extinction coefficients of Cas9, Cyanine dye 3 and Cyanine dye 5.

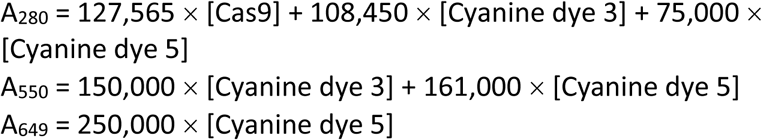 NOTE: The molar extinction coefficient of Cas9 is estimated based on its amino acids sequence using the ProtParam tool in ExPASy^25^. NOTE: The molar extinction coefficient of Cas9 for 550-and 649-nm light and of Cyanine dye 3 for 649-nm light are negligible.
2.7. Snap-freeze the fluorescent Cas9 in 2-5 μL aliquots using liquid nitrogen and store at −80 °C until use. NOTE: This basic method does not separate Cyanine dye 3-Cyanine dye 5 labeled Cas9 from Cyanine dye 3-Cyanine dye 3 and Cyanine dye 5-Cyanince dye 5 labeled Cas9. To separate these, the additional purification step using the hydrophobicity interaction column can be employed^26^.

**Figure 1:**
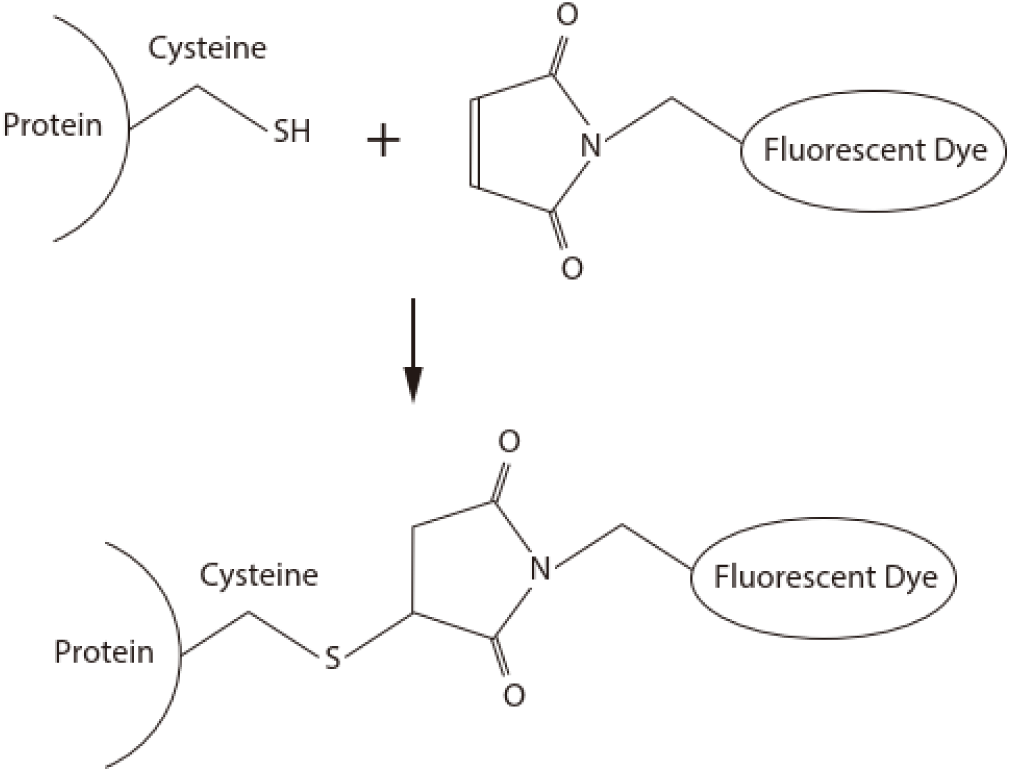
The chemical reaction of maleimide coupling. Maleimide conjugated to a fluorescent dye specifically reacts with the cysteine sulfhydryl group, resulting in covalent binding between the protein and the fluorescent dye.

### Cleavage assay of fluorescent Cas9s

To check the activity of fluorescent Cas9, perform the cleavage assay (Fig. 2). Repeat this cleavage assay for each sample multiple times (at least three times) to ensure the reproducibility of the results.

**Figure 2:**
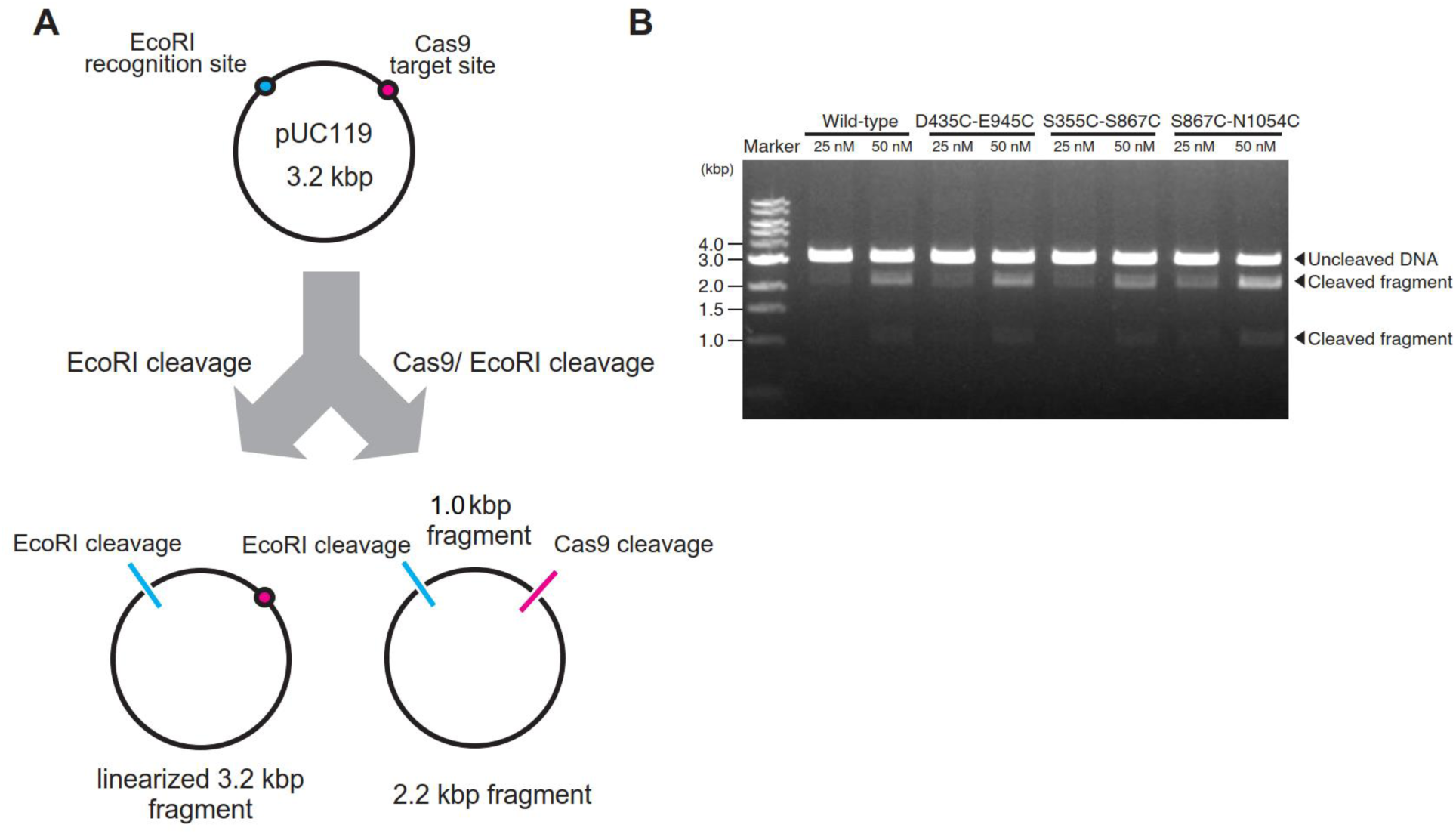
DNA cleavage activity of fluorescent Cas9. **A, Schematic of Cas9 DNA-cleavage assay.** The 3.2 kbp pUC119 plasmid contains both a single EcoRI recognition site and a Cas9 target site. In the presence active Cas9, the plasmid is cleaved into 2.2 kbp and 1 kbp fragments. In the presence of inactive Cas9 protein, the plasmid keeps its original length. **B, DNA cleavage activity of three Cas9 constructs labeled with Cyanine dye 3 and Cyanine dye 5**. After incubating 25 nM or 50 nM Cas9-sgRNA complex, EcoRI, and 5 nM target DNA for 5 min at 37 °C, a fraction of the DNA was cleaved into two fragments. Band intensity analysis suggested that the three FRET constructs retain nuclease activity comparable to that of non-labeled wild-type Cas9 (1.1 ± 0.1 for D435C-E945C, 0.9 ± 0.1 for S355C-S867C, and 1.5 ±0.3 for S867C-N1054C; mean relative activity ± S.E.M. (standard error of the mean), n=3). This figure is reproduced from Ref. 23.

1. Prepare pUC119 plasmid containing a single EcoRI recognition site and a 20-nt target sequence followed by the NGG PAM. NOTE: pUC119 plasmid contains a single EcoRI recognition site at its multi-cloning site.
2. On the day of the experiment, incubate sgRNA at 95 °C for 2 min. After gradual cooling to room temperature, place sgRNA on ice until use.
3. Mix pUC119 plasmid, EcoRI, and Cas9-sgRNA with 10 μL of AB. Incubate the reaction mixture at 37 °C for 5 min.
4. Stop the cleavage reaction by adding a solution of ethylenediaminetetraacetic acid (EDTA; final 40 mM) and Protease K (final 1 mg/mL).
5. Load the reaction mixture on 1% agarose gel. Separate different lengths of DNA by electrophoresis (100 V, 25 min).
6. Soak the agarose gel in liquid containing 0.5 μg/mL ethidium bromide or an alternative fluorescent staining reagent for nucleic acids for 30 min at room temperature. Visualize DNA bands using a UV (ultra violet) or blue-light transilluminator.
7. Capture the image of the DNA bands by camera. Quantify the intensity of each band using the “Gel” analysis tool in ImageJ software^27^.

### Perrin plots to determine orientation factors

To evaluate distance between the two fluorescent dyes on Cas9 from FRET efficiency, each dye should freely rotate during its fluorescence lifetime. Otherwise, orientation between the dyes affects FRET efficiency. To access the rotational mobility of the dyes, the orientation factors are measured using Perrin plots^28^.

1. Prepare 100 μL of 100 nM fluorescent Cas9 (D435C-E945C, S355C-S867C, and S867C-N1054C with no nucleic acid) in cuvettes with 50 μL capacity in buffer (AB + 2.5 mM TSY, 2.5 mM protocatechunic acid (PCA), and 2% protocatechunic acid dioxygenase (PCD)) with or without methyl cellulose (0, 0.001, 0.01, or 0.1%) at room temperature. NOTE: TSY is a commercially available triplet state quencher. Instead of TSY, 1-2 mM 6-hydroxy-2,5,7,8-tetramethylchroman-2-carboxylic acid (Trolox) is also available.
2. Set the parameters of the fluorescence spectrometer (slit width for the emission and excitation, 5 nm; integration time, 1 s; fluorescence and excitation wavelengths for Cyanine dye 3, 566 nm and 554 nm, respectively; fluorescence and excitation wavelengths for Cyanine dye 5, 668 nm and 650 nm, respectively).
3. Measure the fluorescence intensities by manually placing the polarization filters in front of the exciter and detector in the fluorescence spectrometer. Acquire four types of fluorescence intensities:

*I_vh_*, the fluorescence intensity of the horizontal polarization excited by the vertical polarized light;
*I_vv_*, the fluorescence intensity of the vertical polarization excited by the vertical polarized light;
*I_hv_*, the fluorescence intensity of the vertical polarization excited by the horizontal polarized light;
and *I_hh_*, the fluorescence intensity of the horizontal polarization excited by the horizontal polarized light. For better reproducibility, perform three or more measurements in total using independently prepared solutions.
4. Calculate the fluorescence anisotropy, *r*, following the equation described below and using the fluorescence intensities obtained in step 3.

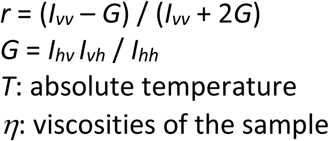 NOTE: *η*values for each methyl cellulose concentration are summarized in Table 1.
5. Plot 1/*r* against *T*/*η*. Calculate the y-intercept (1/*r*_0d_, 1/*r*_0a_) by fitting the plot to a linear function for each fluorescent Cas9 to estimate the anisotropy values. *r*_0d_ and *r*_0a_ are the anisotropy of the donor dye (Cyanine dye 3) and acceptor dye (Cyanine dye 5) in zero viscosity solution, respectively.
6. Using 1/*r*_0d_ and 1/*r*_0a_, calculate *κ*^2^_max_ and *K*^2^_min_ as described below.

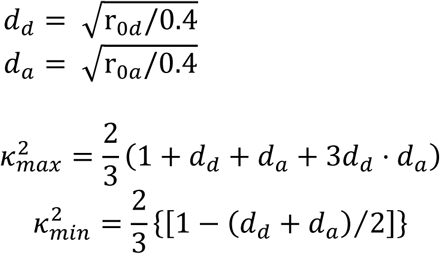

**Table 1:** Viscosity of the buffer against methylcellulose concentrations.

|  |  |  |  |  |
| --- | --- | --- | --- | --- |
| Methyl cellulose concentration | 0% | 0.001% (v/v) | 0.01% (v/v) | 0.1% (v/v) |
| Viscosity (mPa s) | 0.9 | 1.6 | 8.4 | 75.9 |

### Preparation of PEG- and biotin-PEG-coated chambers

Make PEG- and biotin-PEG-coated cover slips for smFRET measurements to avoid non-specific binding of Cas9 to the glass surface^29^ as illustrated in Fig. 3.

**Figure 3:**
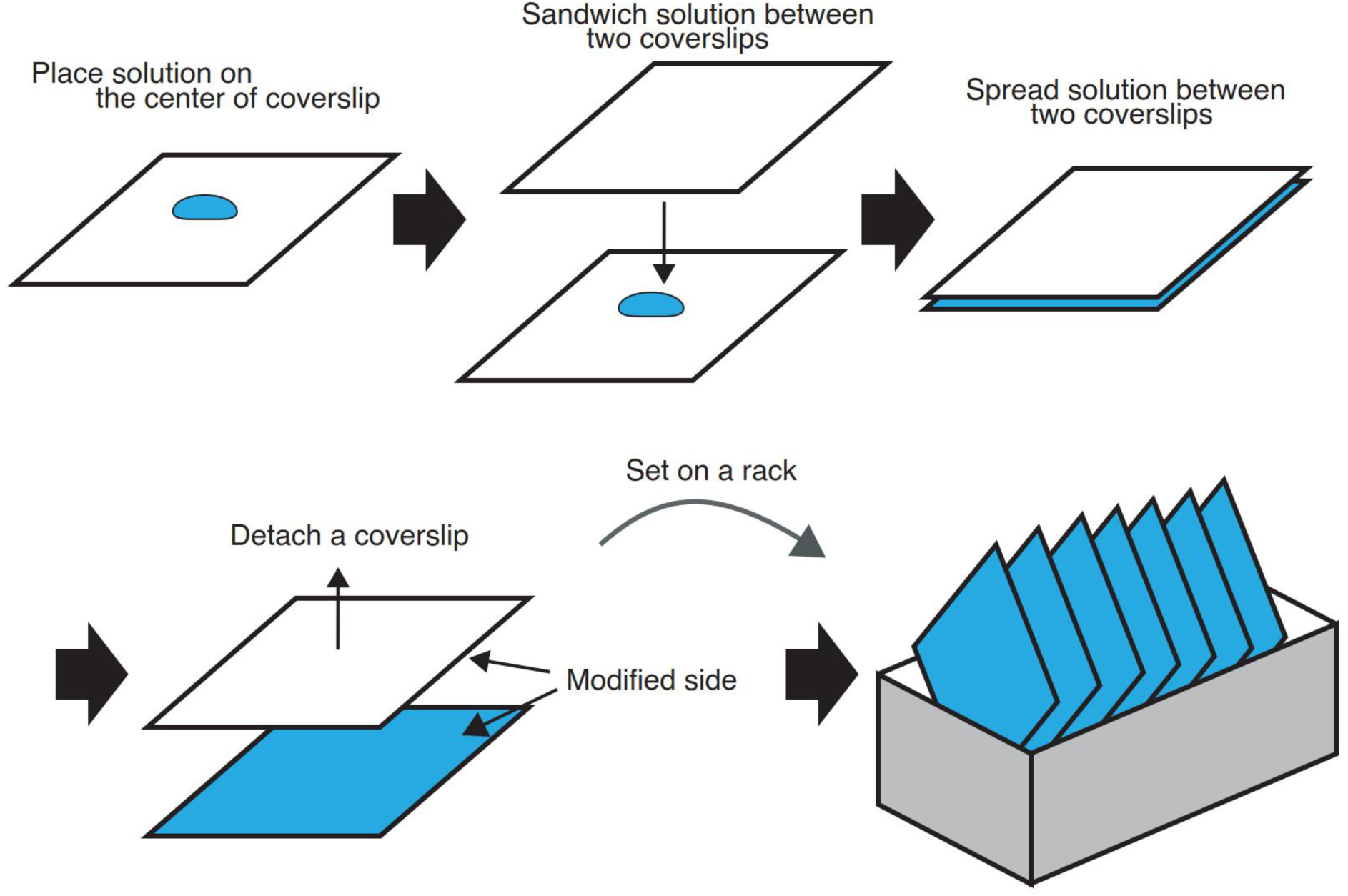
Protocol for modification of glass surface. 10 μL solution containing silane and NHS-PEG or a mixture of NHS-PEG and NHS-biotin-PEG was placed on the center of the cover slip. Next, the solution on the cover slip was sandwiched by another cover slip. Following the spread of the solution between two cover slips, the cover slips were incubated for the time described in the protocol at room temperature. Following the incubation, the sandwiched cover slips were carefully detached using tweezers. Finally, each cover slip was set on a rack and rinsed with ultrapure water.This protocol was applied for both silanization and PEGylation processes except for PEGylation process the incubation was done at a different time in a humid environment.

1. Clean the cover slips (No. 1S, 22 mm × 22 mm).

1.1. Place the cover slips on a rack (typically 10 cover slips). Rinse the cover slips using ultrapure water. Put the rack in a beaker containing 1 N KOH. Clean the cover slips using an ultrasonic washing machine for 20 min.
1.2. Rinse the cover slips with ultrapure water and put the rack in a clean beaker. Pour and exchange ultrapure water in the beaker 20 times to remove KOH completely from the surface of cover slips. Then dry the cover slips in a dryer for 30 min at 80 °C. NOTE: Perform all subsequent procedures in a clean bench.
1.3. Clean the cover slips for 5 min at room temperature using a plasma cleaner and dry them in a dryer.
2. Silanize the cover slips.

2.1. Sandwich 10 µL of N-2 (aminoethyl)-3-aminopropyl-triethoxysilane with two cover slips. Wrap the cover slips using plastic paraffin films.
2.2. Following incubation at room temperature for 20 min, detach the sandwiched cover slips and place the cover slips on a rack. NOTE: DO NOT forget the silanized side of each cover slip.
2.3. Rinse the cover slips on the rack 20 times using ultrapure water and dry them in a dryer for 30 min at 80 °C.
3. PEGylate the silanized side of the cover slips.

3.1. Sandwich 10 µL of 200 mg/mL N-hydroxysuccinimide-polyethylene glycol (NHS-PEG) and 1 mg/mL NHS-PEG-biotin in 50 mM 3-(N-morpholino)propanesulfonic acid (MOPS; pH 7.5) for the observed surface of a flow chamber (8 cover slips in the case of 10 total cover slips) and 200 mg/mL NHS-PEG in 50 mM MOPS (pH 7.5) for the non-observed surface (2 cover slips in the case of 10 total cover slips).
3.2. Following incubation at room temperature for 2 h in a moist environment, detach the sandwiched cover slips and place them on a rack. NOTE: DO NOT forget the PEGylated and biotin-PEGylated side of each cover slip.
3.3. Rinse the cover slips on the rack 20 times with ultrapure water and completely dry them in a dryer.
4. Make 0.5-µL volume micro-chambers (Fig. 4A, B).

4.1. Cut a PEG-coated cover slip into four equal parts (11 mm × 11 mm) using a glass cutter.
4.2. Make three lanes of 1.5 mm width on the PEG-biotin-coated side of 22 × 22 mm cover slip using double-sided adhesive tape (30 µm thickness). Place a PEG-coated small cover slip of 11 × 11 mm, which was cut in the above process, over a PEG-biotin-coated 22 × 22 mm cover slip. NOTE: The PEG-coated side of the small 11 mm × 11 mm cover slip should face the PEG-biotin-coated side of the 22 mm × 22 mm cover slip. Note: A well prepared chamber will show no significant leakage (Fig. 4C). Additionally, following wash-out of the colored solution, the remaining dye in the solution is not visible (Fig. 4D). These observations suggest that the micro-chamber made with double-sided adhesive tape is adequate for the following smFRET measurements.
4.3. Check the cleanliness of the micro-chamber and surface passivation by fluorescence microscopy. NOTE: The micro-chamber will show some fluorescent spots (Fig. 5A). The biotinylated sample, such as biotinylated Cas9, specifically binds to the glass surface via avidin-biotin binding (Fig. 5B).
4.4. Individually vacuum seal the micro-chambers and store them at -80 °C until use.

**Figure 4:**
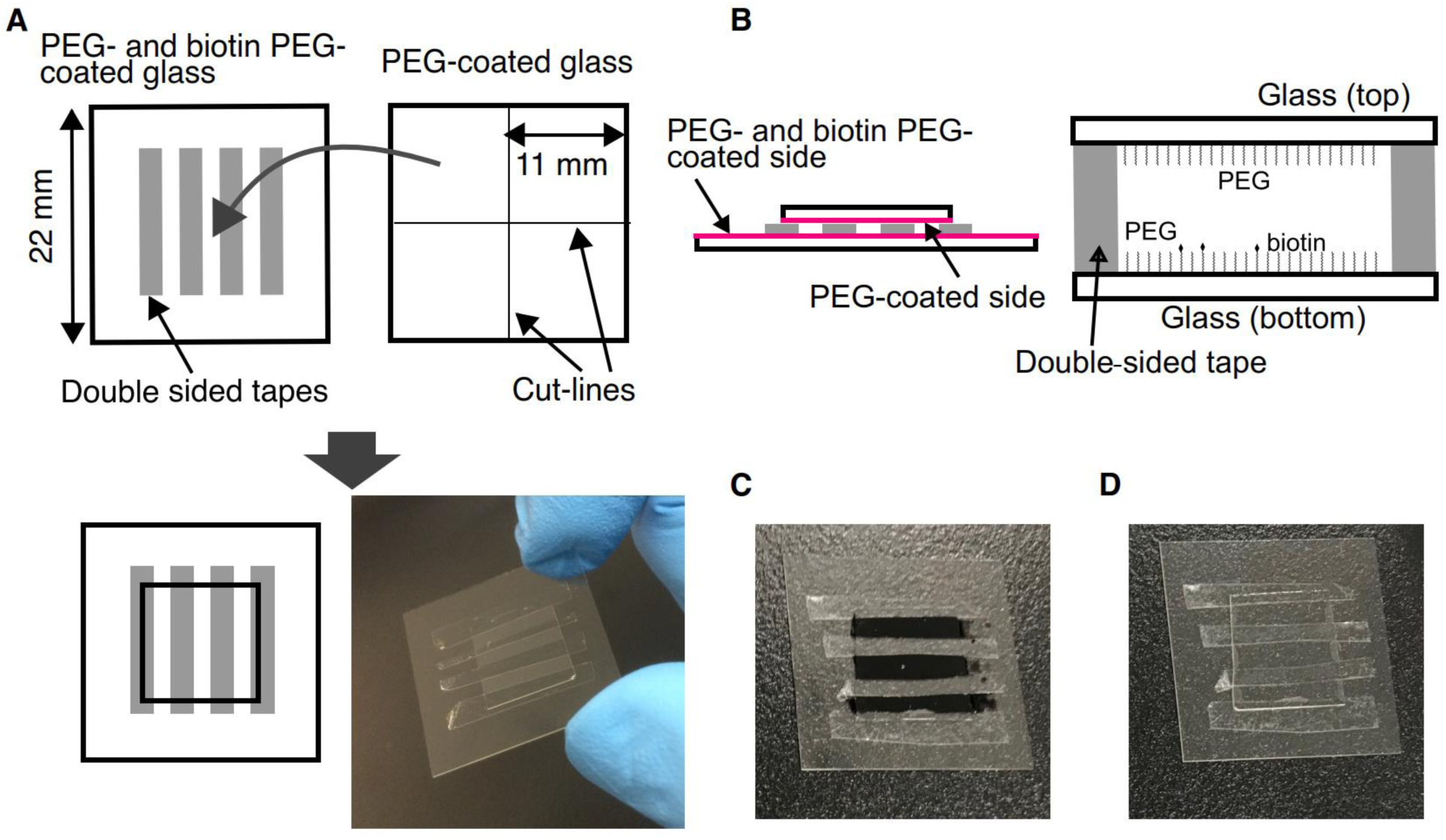
Preparation of the glass chamber. **A, Schematic of the glass chamber preparation.** Among 10 washed cover slips, 2 cover slips were coated with PEG and the other 8 cover slips were coated with PEG and PEG-biotin. Each of the 2 PEG-coated cover slips were cut into four pieces. The glass chamber was made by adhering the small PEG-coated and the large PEG-biotin-coated cover slips using 30-μm thick double-faced tape. **B, PEG- and PEG-biotin-coated side of the cover slips.** To construct the glass chamber, the coated side of the cover slips is shown in magenta. The top cover slip is coated with PEG, and the bottom one is coated with PEG and PEG-biotin. **C, Photograph of the glass chambers filled with dye solution.** The three lanes were filled with a solution of black dye. **D**, Following three washes with 2 µL of ultrapure water, the black dye solution in the chamber was replaced with transparent ultrapure water.

**Figure 5:**
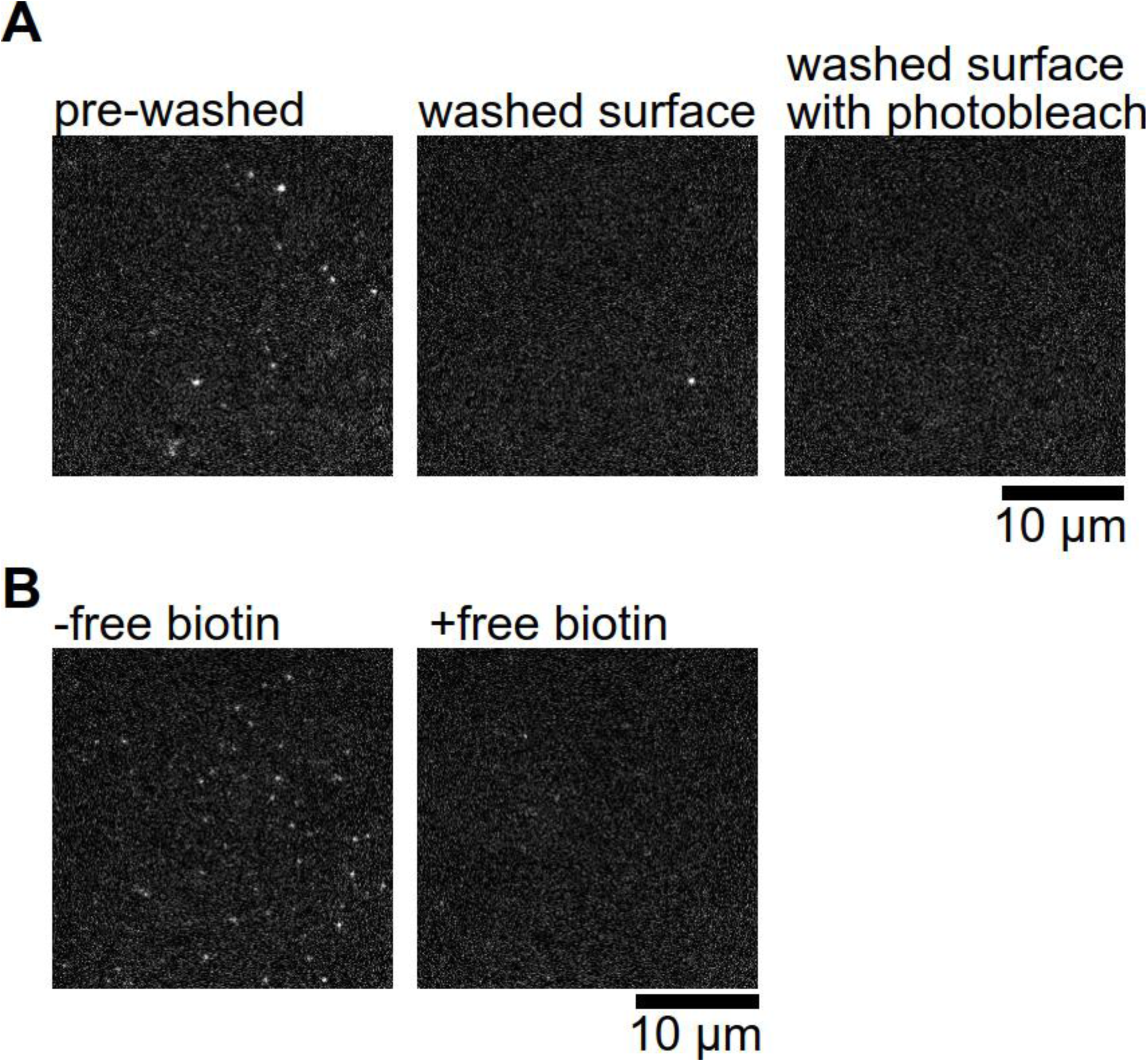
Cleanliness and passivation of the glass surface. **A, Fluorescent images of the glass surface with and without cleaning.** Many fluorescent artefacts were observed before cleaning (left panel), but only few of them remained after cleaning (middle panel). The remaining artefacts were photobleached before introducing the Cas9 molecules into the chamber (right panel). **B, Confirmation of surface passivation by PEG.** Biotin-fluorescent Cas9 molecules (bright spots) were anchored on the PEGylated surface through biotin-PEG and neutralized avidin (left panel). However, when biotin binding sites in neutralized avidin were blocked by free biotin, the biotin-fluorescent Cas9 molecules did not bind to the surface, suggesting that PEG effectively suppresses non-specific binding on the surface.

### Single-molecule FRET measurements under total internal reflection fluorescence microscopy (TIRFM)

1. Load 1 µL of 0.1-1 mg/mL neutralized avidin in AB into the micro-chambers prepared in step 4 of the previous section (Fig. 6A).
2. Following incubation for 2 min at room temperature, remove excess neutralized avidin by three washes with 2 µL AB. Note: Exchange liquid in the chamber by adding fresh liquid from an open end of the chamber, while using a piece of sliced filter paper at the other end as an absorber (Fig. 6A).
3. Place the chamber on the stage and fix the corners of the chamber on the stage using tape. Put a small cap with wet paper on the micro-chamber to keep a humid environment (Fig. 6B).
4. Illuminate the glass surface in the micro-chamber with a 532-nm laser for 40 s per one image field using fluorescence microscopy to photobleach residual fluorescent particles on the glass surface. NOTE: Check “Multiple positions (XY)” in the “Multi-Dimensional Acquisition” panel and record the xy-stage position by pushing the button “Edit position list”. During the xy-stage position recording, only the XY position should be checked (the other options such as Z and autofocus should be unchecked). The acquisition order should be “Time, position”. After completion of the above settings, push the button “Acquire!” to initiate sequential image acquisitions with the 532-nm laser (“Auto shutter” in the main panel should be checked) using open source microscopy software (Micro-Manager, Open Imaging)^30^. The laser power is set to maximum power level for this procedure. NOTE: DO NOT move the chamber on the stage in the following procedure in order to acquire fluorescent images in the same area.
5. In the meantime, incubate 0.3-1 nM fluorescent Cas9 with 200 nM single guide RNA (sgRNA) for 2 min at room temperature in a 0.6-mL tube for sgRNA-bound fluorescent Cas9 imaging, or with 200 nM sgRNA and 200 nM plasmid DNA containing the Cas9 cleavage site for sgRNA-and DNA-bound Cas9 imaging. NOTE: Prior to mixing sgRNA and Cas9, heat the sgRNA at 95 °C for 2 min. Following gradual cooling to room temperature, place the sgRNA on ice until use. This heating and cooling process should be done on the day of use.
6. Load 2 µL of 0.3-1 nM biotinylated fluorescent Cas9 into the micro-chamber. Note: As an alternative method, immobilize fluorescent Cas9 on the glass surface via biotinylated nucleic acid (sgRNA or DNA). In this case, FRET efficiency of the Cas9 is measurable only when the Cas9 molecule binds with biotinylated nucleic acid.
7. Following incubation of the biotinylated fluorescent Cas9 for 2 min at room temperature, remove excess fluorescent Cas9 with three washes of 2 µL AB. NOTE: Fluorescent Cas9 molecules specifically adsorb onto the glass surface via avidin-biotin interactions. When 0.3 nM fluorescent Cas9 was applied to the chamber, the fluorescent Cas9 density on the glass surface is 33.6 ± 5.5 × 10^−3^ fluorescent spots/µm^2^ [n = 6 fields; mean ± s.d. (standard deviation)], resulting in a microscopic field of about 55 µm2 with few overlapping particles but a sufficient number of fluorescent spots (~100).
8. Load 2 µL AB with 2.5 mM TSY, 2.5 mM PCA, and 2% PCD solution into the micro-chamber. NOTE: Add 200 nM sgRNA to the buffer for sgRNA-bound fluorescent Cas9 imaging, or 200 nM sgRNA and 200 nM plasmid DNA for sgRNA- and DNA-bound Cas9 imaging.
9. Put a small cap with wet paper on the micro-chamber to keep a humid environment (Fig. 6B). Wait 5 min to avoid any large xy drift of the stage (Fig. 7). Alternatively, correct drift of the image before analysis.
10. Set the acquisition parameter in the microscopy software. Acquire the fluorescence image. NOTE: Set the EMCCD in frame-transfer mode and select the laser power values (typically 12.5 and 5 mW for 532-nm and 642-nm lasers, respectively). Data acquisition is performed with an acquisition rate of 10 frames/s and total frame number of 1,201 for D435C-E945C and S355C-S867C and 401 for S867C-N1054C using Micro-Manager^30^. Illuminate the same field with the 642-nm laser to directly excite the Cy5 fluorescence prior and following the smFRET measurement to count double-labeled Cyanine dye 3 and Cyanine dye 5 molecules and distinguish the termination of FRET from photobleaching. NOTE: Repeat at least three observations for each condition using different chambers to confirm reproducibility.

**Figure 6:**
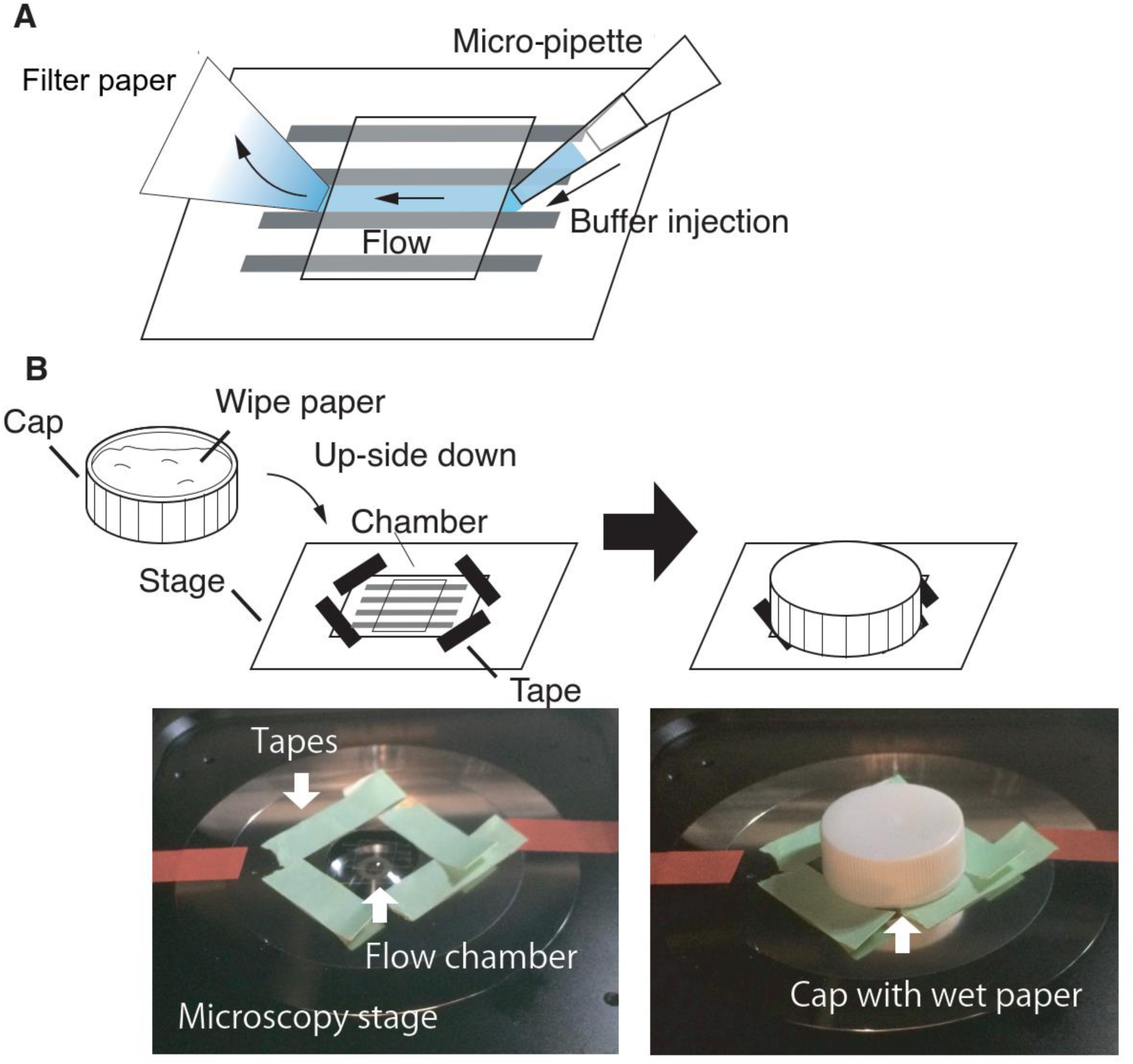
Experimental procedures. A, Schematic drawing of buffer exchange. To exchange the buffer solution, new buffer (4 times the volume of the chamber capacity; i.e., 2 µL buffer for a 0.5 µL chamber) is injected from one side of the chamber using a micro-pipette, while the buffer existing in the chamber was absorbed by filter paper on the other side. **B, Schematic drawing and photograph of the experimental setup.** The four corners of the microchamber were fixed on the microscope stage plate with four strips of tape (green). The microscope stage plate was also fixed to the stage with at least two strips of tape (red) to prevent rotation of the plate on the stage. A small cap (typically the cap of a centrifuge tube) was stuffed with wet paper and placed on the microchamber fixed on the microscope stage to keep the microenvironment humid.

**Figure 7:**
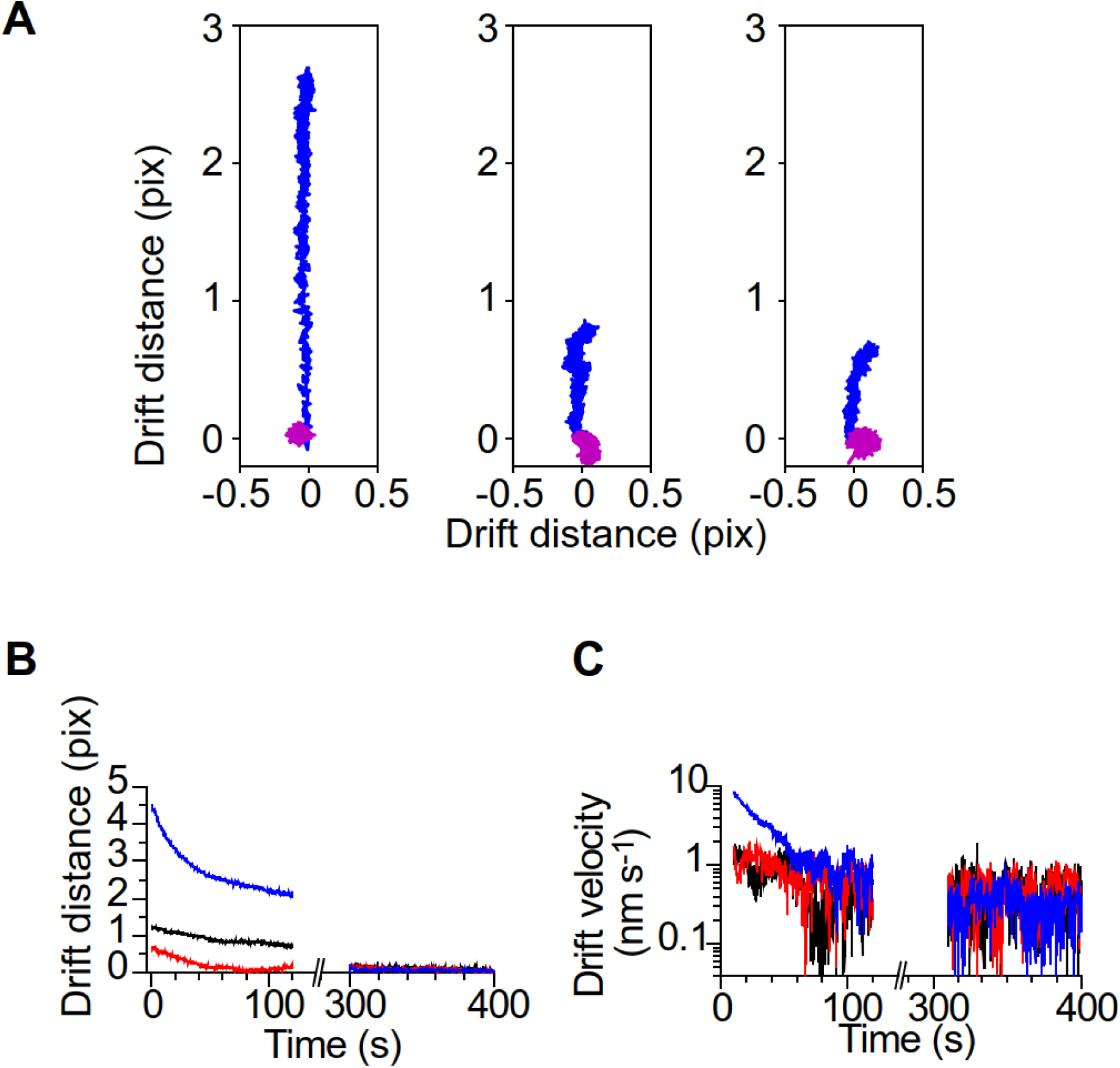
Drift of the microscope stage. A, Time trajectories of fluorescent beads adsorbed on the glass surface. Time trajectories of fluorescent beads illuminated with a 532-nm laser were tracked for 2 min immediately after the introduction of 2 µL AB into the chamber (blue) and 5 min after the introduction (magenta). Centroid positions of the fluorescent beads were calculated using ImageJ. The initial point for each tracking was set to [0, 0]. **B, Drift distances of the beads.** Three representative data sets show large stage drift in the first 2 min after the introduction of AB. After 5 min, the large drift (~1 pixel per 100 s) was not observed (blue: left trajectory in A; black: middle trajectory in A, red: right trajectory in A). **C, Velocity of the stage drift.** In the first ~50 s, the velocity was faster than 1 nm s-1, which corresponds to 1-pixel movement per 108 s. The color difference is the same as in B.

### Data analysis of single-molecule FRET

1. Load the time series of fluorescent images (tiff file sequence saved using Micro-Manager) using the custom program written in Python.
2. Find Cyanine dye 3- and Cyanine dye 5-labeled fluorescent particles and make the region-of-interest (ROI) with 6-pixel width around the centroid of the fluorescent particle.
3. Convert the time series of the fluorescent images within each ROI into time trajectories of fluorescence intensities (*I_obs_A_* and *I_obs_D_* for the acceptor and donor, respectively).
4. Calculate the background intensity based on the mean intensity in each ROI after the photobleaching of fluorescent Cas9 (*I_back_A_* and *I_back_D_* for the acceptor and donor, respectively). Subtract the background intensity from the observed fluorescence intensity.

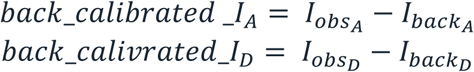

To estimate accurate fluorescence intensities of the donor and acceptor, calibrate ratios of measured intensities of each dye in the different fluorescence channels. This ratio is called leakage ratio as it corresponds to leakage of fluorescence in undesired channel. Since the ratio highly depends on the utilized microscopy system (in particular, the filter set of the emission filter and dichroic mirror), the ratio should be estimated for each system used. In the below equation, *leak_calibrated_I_A_* and *leak_calibrated_I_D_* are the background and leakage-calibrated fluorescence intensity of the acceptor and donor, respectively. *r_DA_* and *r_AD_* are the leakage ratio of the donor fluorescence in the acceptor fluorescence channel and the leakage-ratio of the acceptor fluorescence in the donor fluorescence channel, respectively.

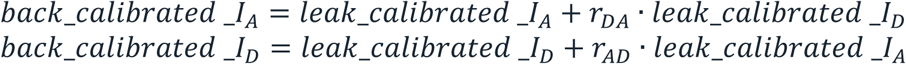
5. Calculate the time trajectory of FRET efficiency using the equation based on the calibrated fluorescence intensities of Cyanine dye 3 and Cyanine dye 5.

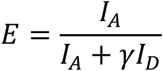

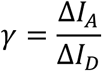 NOTE: *I_A_* is the calibrated fluorescence intensity of Cyanine dye 5 (acceptor) and *I_D_* is that of Cyanine dye 3 (donor). γ is the correction factor, which accounts for differences in the quantum yield and detection yield derived from the sensitivity of the camera and the properties of the dichroic mirrors and emission filter.
6. To determine the time of the state transitions in each smFRET time trajectory, perform the analysis based on a hidden-markov-model (HMM)^24^. NOTE: The hmmlearn package for Python^31^ and HaMMy^24^ are two good examples of a program for the HMM analysis.
7. Plot the transition density plot (TDP) using a graph visualization tool such as the matplotlib library for Python^32^ to visualize the relationship between successive transitions in the states based on FRET efficiency^23^ (Fig. 8).
8. To quantify state transitions based on TDP, classify the density into groups using the k-means clustering method. Choose the optimal number of groups using Silhouette analysis (Fig. 8). NOTE: There are many analytical tools for k-means clustering^33^. The scikit-learn package in Python^34^, for instance, can be utilized.

**Figure 8:**
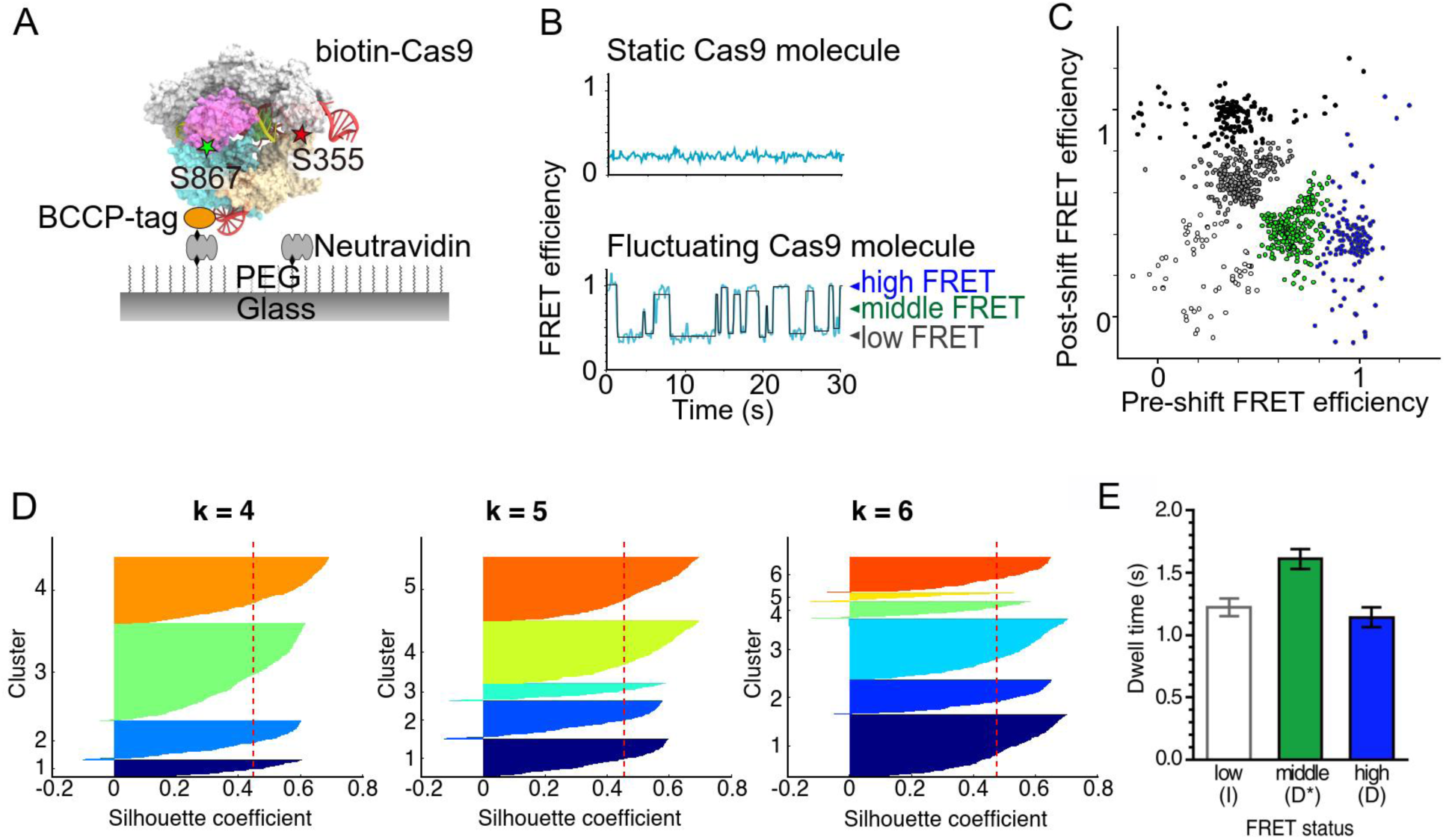
Inter-domain flexibility in the sgRNA-DNA-Cas9 complex. A, Schematic of the smFRET system. The sgRNA-DNA-Cas9 ternary complex was anchored on a PEG- and biotin-PEG-coated glass surface via neutralized avidin-biotin linkage. **B, Representative time trajectory of S355C-S867C (HNH-REC) Cas9 construct.** ~80% of the Cas9 molecules showed stable low FRET efficiency (upper panel), while the rest of the molecules demonstrated fluctuations (bottom panel). HMM analysis (black line) suggests that the fluctuating molecules adopt three FRET states. **C, Transition density plot of FRET efficiency fluctuations.** The k-means analysis classified the plots into five groups, suggesting that direct transition between the high and middle FRET states is rare. This figure is reproduced from Ref. 21 with some modifications. **D, Silhouette plots.** The Silhouette coefficients, which were calculated using the machine learning Python Package Scikit learn, are plotted for each cluster in the cases of k = 4, 5, and 6. The vertical red dashed lines indicate the mean values of the Silhouette coefficients. In the cases of k = 4 and 5, all clusters showed Silhouette coefficients higher than the mean values. This was not true for k = 6, meaning that k = 5 is the most probable number of clusters for the transition density plot shown in C. **E, Bar plot of dwell times for each transition.** The mean dwell times were determined by fitting the dwell time distributions (n = 399, 223, and 136 for low, middle, and high FRET states, respectively) to a single exponential decay function. Error bars show SEM. This figure is reproduced from Ref. 23.

## REPRESENTATIVE RESULTS

smFRET measurements require a protein be labeled with two distinct fluorochromes and retain its normal activity. In the present protocol, two cysteine residues were introduced into Cas9 by genetic modification and labeled with Cyanine dye 3- and Cyanine dye 5-maleimide. Typical labeling efficiency of each dye on Cas9 was 60-80%, indicating that approximately 40-60% of Cas9 molecules were labeled with both dyes. The low labeling efficiency makes it difficult to find two-colored Cas9 molecules. In these cases, we repeated the labeling procedure. The main causes of the low labeling efficiency are the pH of the reaction mixture and the deterioration of maleimide. The maleimide group reacts specifically with sulfhydryl groups between pH 6.5 and 7.5. It also reacts with primary amines and hydrolyzes into non-reactive maleamic acid at pH over 8.5. Thus, keeping maleimide active by avoiding hydrolysis is crucial for efficient labeling of the protein.

We also confirmed the DNA-cleavage activity of fluorescent Cas9. Approximately 3.2 kbp of pUC119 plasmid was linearized by EcoRI restriction enzyme. In the presence of the sgRNA-Cas9 binary complex with Mg^2+^ ion, the linearized DNA was cleaved into 2.2 kbp and 1 kpb fragments (Fig. 2A). From the intensity of the bands, we estimated the DNA-cleavage activity of each Cas9 protein. All fluorescent Cas9 showed almost the same level of DNA-cleavage activity (>90%) as non-labeled wild-type Cas9 (Fig. 2B). The retained activity confirmed the feasibility of fluorescent Cas9 for the following smFRET measurements.

For accurate smFRET measurements, the cleanliness and passivation of the glass surface on the observatory chamber are important. The chamber should be prepared under clean conditions using a general clean booth or biological clean hood of at least class 1,000 to class 10,000. Otherwise, fluorescent artefacts are observed even on the washed glass surface. Even with ideal cleaning and passivation conditions, sometimes fluorescent artefacts remained on the surface. In these cases, the artefacts were photobleached before the smFRET observation (Fig. 5A), allowing us to distinguish the signals from fluorescent Cas9 molecules. The procedures of the neutralized avidin and Cas9 anchoring on the glass surface did not require a clean booth. With filtered solutions, fluorescent artefacts did not increase significantly upon the introduction of solutions into the glass chamber in a standard laboratory (Fig. 5A). Our method successfully produced a clean glass surface with anchored Cas9 molecules.

To avoid protein inactivation caused by non-specific binding to the glass surface, it is necessary to passivate the surface. We coated the surface with polyethylene glycol (PEG), which is widely used for passivation of a glass surface. We mixed biotin-PEG with intact trimethyl-PEG. The biotinylated Cas9 was anchored on the surface via avidin-biotin linkage. When the biotin binding sites in neutralized avidin on the surface were blocked by excess free biotin, biotin-labeled fluorescent Cas9 did not bind to the surface (Fig. 5B). This result suggests that PEG effectively prevents non-specific binding on the surface, so that Cas9 molecules are specifically anchored on the surface through their biotin moiety.

Following the above precautions, we performed smFRET measurements under TIRFM. Through a color splitter, fluorescence from Cyanine dye 3 and Cyanine dye 5 were simultaneously recorded on distinct positions of an electron-multiplier CCD camera. We found that the recorded position of the Cas9 molecules moved 100-300 nm within 100 s immediately after the solution exchange due to the drift of the microscope stage (Fig. 7). This movement corresponded to 1-3 pixels of the recorded movie; thus, image drift correction should be employed to track and analyze Cas9 molecules. Alternatively, we started the recording 5 min after introducing the final solution into the chamber. In this case, the stage drift was suppressed within 0.3 pixels during 120-s observations (Fig. 7). Because the fluorescence intensity in 6×6 pixel ROI was used to estimate the fluorescence intensity derived from fluorescent Cas9, stage drift did not affect the results in our observation time (120 s). Therefore, the movies taken with this method were analyzed without drift correction. Furthermore, we concluded the solution drift inside the chamber was also negligible, because we detected specific FRET efficiency distributions including a static FRET efficiency by using three different FRET Cas9 constructs (D435C-E945C, S355C-S867C, and S867C-N1054C) in four different nucleic acid-binding conditions (Apo, sgRNA binding, sgRNA/target DNA binding and sgRNA/target DNA/Mg^2+^ binding).

In our assay, 68-96% of double-labeled Cas9 molecules showed anti-correlated changes in the fluorescence intensities of Cyanine dye 3 and Cyanine dye 5, which indicates FRET between the two dyes. The FRET efficiency was calculated from the intensities until photobleaching of one of the dyes. Among the molecules showed anti-correlated intensities of the two dyes, ~80% of the sgRNA-DNA-Cas9 ternary complexes showed relatively constant FRET efficiencies during the observation period. Contrarily, the other ~20% of complexes showed fluctuations, suggesting flexible inter-domain movements in the molecules (Fig. 8B). HMM and k-means analyses of the FRET efficiency trajectories suggested that the HNH domain can translocate its position against the REC lobe to adopt three conformations (Fig. 8). The highest FRET efficiency state only appeared in the fluctuating complex. As the appearance of this state closely correlate the DNA cleavage activity of Cas9, our results suggest that movements in the flexible domain of the Cas9 protein allow Cas9 to take a transient conformation that is responsible for cleavage of the target DNA.

To interpret smFRET data, the orientation factor, *K*^2^, should be calculated from the fluorescence anisotropy data, because *K*^2^ determines the contribution of the orientation of the two dyes in FRET efficiency. When the two dyes freely rotate during their fluorescence lifetime, *K*^2^ becomes 2/3 and the FRET efficiency directly reflects the change in distance between the two dyes. However, Cyanine dye 3 and Cyanine dye 5 on Cas9 showed high (~0.3) anisotropy, meaning *K*^2^ is widely distributed (in the range of 0-4). In this case, the FRET efficiency is affected by both the distance and the orientation between the dyes. Here, the Förster distance (R_0_) is described using the below equation.

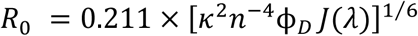

(*n*: refractive index of the medium; *φ_D_*: fluorescence quantum yield of the donor; *J(λ)*: spectral overlap between the donor and the acceptor)

The widely distributed *K*^2^ for the Cyanine dye 3-Cyanine dye 5 pair on Cas9 obscure the value of R_0_, resulting in ambiguous distances between the dyes based on the FRET efficiency according to the equation:

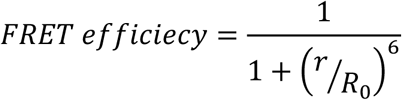

(*r*: distance between donor and acceptor).

However, high anisotropy is not necessarily undesirable, because conformational changes coupled with not the distance but with the orientation of the dyes can be detected. In the case that low anisotropy is preferred, another combination of fluorescent dyes with long fluorescence lifetimes is recommended, because Cyanine dye 3 and Cyanine dye 5 show relatively short lifetimes (< 1 ns)^23^.

## DISCUSSION

Here we described a detailed procedure for Cas9 smFRET measurements. One of the critical points of the measurement is to keep Cas9 molecules active after fluorescent labeling and anchoring them to the glass surface. For fluorescent labeling, cysteine-maleimide coupling is widely used. Our fluorescently labeled Cas9 retained robust DNA cleavage activity, but sometimes the mutagenesis and labeling of the cysteine residues compromise protein activity. In these cases, using other residues for cysteine-substitution or site-specific labeling methods, such as click chemistry with unnatural amino acids, should be employed. As for anchoring on the glass surface, passivation of the surface is essential, because non-specific hydrophobic interactions between a protein and the surface could change the protein conformation. Note that the appropriate reagents for passivation depend on the observed bio-molecules and the surface conditions. For instance, casein is necessary for the single-molecule observation of microtubule-based motor movement^35,36^, but not in the case of Cas9 conformational changes^23, 37, 38^. It is always necessary to confirm that the observed protein has anchored on the surface via the intended linker.

Our smFRET measurements detected the conformational dynamics of Cas9. Visualizing the conformational dynamics is unique feature of smFRET compared to crystal and cryo-electron microscopy structural analyses, which capture a snapshot of the conformation. Recent progress on high-speed atomic force microscopy has enabled to visualize the conformational dynamics of Cas9 during target DNA binding and cleavage while also capturing the whole structure of the complex^39^. This indeed is a big advantage of the atomic force microscopy, considering smFRET can only detect the dynamics of fluorescently labeled positions. Yet, smFRET has higher spatial and temporal resolutions than atomic force microscopy when monitoring two positions with distance close to a Förster distance of florescence dyes (typically ~5 nm). Moreover, smFRET data combined with molecular dynamics simulations can produce a complementary model for conformational dynamics of whole domains in a protein^40^, suggesting that this combination is a prominent tool for studying nanoscale conformational dynamics.

The application of smFRET to study the dynamics of gene-editing proteins like Cas9 and Cas13 is improving our ability to design gene-editing proteins of higher efficiency such as hyper-accurate Cas9 variant (HypaCas9)^41^. Furthermore, by utilizing fluorescent DNA targets, smFRET have revealed how Cas9 distinguishes on-target and off-target DNA molecules^42–44^. Overall, the revelation of protein dynamics through smFRET is expected to significantly advance the design of better gene-editing tools.

## ACKNOWLEDGMENTS

This study is supported by JST, CREST, JPMJCR14W1 (to S.U), MEXT, Grand-in-Aid for Scientific Research (C), 18K06147 (to T.S.), and MEXT, Grants-in-Aid for Young Scientists (B), 15K18514 (to T.S.). We thank members of the Uemura and Nureki laboratories for valuable discussions. We also thank P. Karagiannis (Sofia Science) for helpful discussions and comments on the manuscript.

